# The role of GpsB in cell morphogenesis of *Staphylococcus aureus*

**DOI:** 10.1101/2023.06.16.545294

**Authors:** Sara F. Costa, Bruno M. Saraiva, Helena Veiga, Leonor B. Marques, Simon Schäper, Marta Sporniak, Daniel E. Vega, Ana M. Jorge, Andreia M. Duarte, António D. Brito, Andreia C. Tavares, Patricia Reed, Mariana G. Pinho

## Abstract

For decades, cells of the gram-positive bacterial pathogen *Staphylococcus aureus* were thought to lack a dedicated elongation machinery. However, *S. aureus* cells were recently shown to elongate before division, in a process that requires a SEDS (Shape Elongation Division and Sporulation) / PBP (Penicillin Binding Protein) pair for peptidoglycan synthesis, consisting of the glycosyltransferase RodA and the transpeptidase PBP3. In ovococci and rod-shaped bacteria the elongation machinery, known as elongasome, is composed of various proteins besides a dedicated SEDS/PBP pair. To identify proteins involved in the elongation of *S. aureus*, we screened the Nebraska Transposon Mutant Library, which contains transposon mutants in virtually all non-essential staphylococcal genes, for mutants with modified cell shape. We confirmed the roles of RodA/PBP3 in *S. aureus* elongation and identified GpsB, SsaA, and RodZ as additional proteins involved in this process. The *gpsB* mutant showed the strongest phenotype, mediated by the partial delocalization from the division septum of PBP2, the only bifunctional PBP in *S. aureus*, with both glycosyltransferase and transpeptidase activity, and of the PBP4 transpeptidase. Increased levels of these PBPs at the cell periphery result in higher levels of peptidoglycan insertion throughout the entire cell, overriding the RodA/PBP3-mediated peptidoglycan synthesis at the outer edge of the septum, which leads to cell elongation. As a consequence, in the absence of GpsB, *S. aureus* cells become more spherical.

## Introduction

*Staphylococcus aureus* is a gram-positive bacterial pathogen, responsible for a high number of antibiotic-resistant infections, and is currently the second major cause of death by antibiotic-resistant infections (1). For decades, *S. aureus* cells were thought to lack a dedicated elongation machinery and *S. aureus* was often mentioned as a canonical example of truly spherical cocci. Rods, which have a cylindrical shape, undergo cell elongation before the synthesis of a division septum, followed by division of the mother cell into two identical daughter cells. These two processes, cell elongation and division, are organized by two cytoskeletal proteins, the bacterial actin homologue MreB, and the tubulin homologue FtsZ, respectively, which coordinate the assembly of two multiprotein complexes, the elongasome or Rod complex and the divisome (2-4). These complexes include cell wall synthesis proteins and organize synthesis of new peptidoglycan at the lateral wall and at the division septum of rods.

*S. aureus* lacks an MreB homologue and was previously thought to lack a canonical elongasome (5). The view that *S. aureus* had only one cell wall synthesis machinery that made peptidoglycan at the division septum, was present in work by G. Satta in the 1990s (6). This work proposed a “two-competing-sites model” for peptidoglycan assembly which predicted that there were two types of cocci: one that synthesized peptidoglycan only at the septum (which would include *S. aureus*) and another that also synthesized peptidoglycan for lateral wall elongation (which would include ovococci such as *Enterococcus faecium* or *Streptococcus agalactiae*) (6). In these studies, *S. aureus* was unable to elongate under various conditions that promoted elongation of ovococci, such as the presence of antibiotics that inhibit septation (6). However, the recent development of super-resolution microscopy techniques allowed imaging of the small (1μm in diameter) staphylococcal cells with sufficient resolution to observe minor morphological changes that occur during the cell cycle. This showed that *S. aureus* cells do undergo slight elongation, reflected in an increase of the ratio of the longer cell axis (perpendicular to the division septum) to the shorter cell axis (overlapping the septum) during the cell cycle (7). The *S. aureus* cell cycle can be divided into three different phases: Phase 1, before initiation of division septum synthesis, Phase 2, during which septum synthesis occurs, and Phase 3, during which cells have a complete septum, undergoing maturation, before it is split into two identical daughter cells (7). Cell volume increases during the entire cell cycle, while cell elongation occurs mostly during Phases 1 and 3 (7). We have recently shown that this elongation process is dependent on the presence of the SEDS (Shape Elongation Division and Sporulation) / PBP (Penicillin Binding Protein) pair RodA/PBP3, proteins with glycosyltransferase activity responsible for glycan strand synthesis, and transpeptidase activity responsible for peptidoglycan crosslinking, respectively (8). Together these proteins catalyse peptidoglycan synthesis, including at the sidewall of the septal region, resulting in slight cell elongation (8). This midcell synthesis is reminiscent to that observed in ovoccocoid bacteria such as *Streptococcus pneumoniae*, that also lack an MreB homologue (9, 10), but different from elongation of rod-shaped bacteria like *Escherichia coli* or *Bacillus subtilis*. In the latter, MreB polymerizes into short filaments that move processively in a circumferential direction, perpendicular to the long axis of the rod, organizing the elongasome machinery, and leading to peptidoglycan incorporation over the entire length of the cell (11-13). Besides having a dedicated SEDS/PBP pair, rods elongasome also includes two transmembrane proteins MreC and MreD, proposed to couple intracellular MreB to extracellular PBPs (14) or RodZ, which interacts with other elongasome components (15) and may modulate MreB filament density (16).

Given the relative complexity of elongasomes in rods, it seemed unlikely that elongation in *S. aureus* would require only RodA and PBP3, the only two proteins so far identified as having a role in this process. We hypothesized that *S. aureus* elongation was not essential during growth in rich medium, as deletion of both *rodA* and *pbp3* genes was not lethal in these conditions (8). Therefore, to identify additional factors required for *S. aureus* elongation we screened by microscopy the Nebraska Transposon Mutant Library (NTML), composed of mutants with a *bursa aurealis* transposon insertion in virtually every non-essential gene of the methicillin resistant *S. aureus* (MRSA) strain JE2 (17), for mutants with cells with decreased eccentricity.

## Methods

### Bacterial growth conditions

*S. aureus* strains were grown at 37 °C in tryptic soy broth (TSB, Difco) with agitation, or on plates of tryptic soy agar (TSA, Difco). When necessary, culture media was supplemented with appropriate antibiotics (chloramphenicol 10 μg/mL, erythromycin 10 μg/mL (Sigma-Aldrich, deletion mutants) or erythromycin 25 μg/mL (NTML mutants), kanamycin 150 μg/mL (Apollo Scientific, overnight cultures) or 50 μg/mL (cultures for microscopy), or a combination of kanamycin (50 μg/mL) with neomycin (50 μg/mL, Apollo Scientific), with 100 μg/mL 5-bromo-4-chloro-3-indolyl-β-D-galactopyranoside (X-Gal, VWR), or with 0.1 μM CdCl_2_ (Sigma-Aldrich). *E. coli* strains were grown at 37 °C in Luria-Bertani broth (LB, Difco) or on LB agar (Difco) supplemented with ampicillin (100 μg/mL, Sigma-Aldrich) when required.

### Microscopy screening of the Nebraska Transposon Mutant Library (NTML)

Batches of 48 mutants of the NTML (17) were imaged, with JE2 imaged before and after the first and last group of 8 mutants, respectively. Each strain was grown overnight at 37 ºC in TSB supplemented with erythromycin 25 μg/mL, with the exception of JE2 which was grown without antibiotic. Each overnight culture was back-diluted 1:200 in TSB and grown until OD_600nm_ 0.8. From each culture, a 300 μL aliquot was incubated with 0.4 μL of both Nile red (10 mg/mL, Invitrogen) and Hoechst 33342 (5 mg/mL, Invitrogen) for 5 minutes at 37 ºC with shaking. Samples were pelleted and resuspended in 10 μL of a solution of 1:3 (vol/vol) TSB/ phosphate buffered saline (PBS, NaCl 137 mM, KCl 2.7 mM, Na_2_HPO_4_ 10 mM, KH_2_PO_4_1.8 mM). One microliter of each sample was mounted on a layer of 1.2% (w/v) agarose in 1:3 (vol/vol) TSB/PBS placed on a glass plate (Bio-Rad Mini-PROTEAN Short Plate) with a coverslip placed on top of each sample. Eight samples were imaged per glass plate, using a Zeiss Axio Observer microscope equipped with a Plan-Apochromat 100x/1.4 oil Ph3 objective, a Retiga R1 CCD camera (QImaging), a white-light source HXP 120 V (Zeiss) and Metamorph 7.5 software (Molecular Devices). The filters (Semrock) Brightline TXRED-4040B (Nile red) and Brightline DAPI-1160A (Hoechst 33342) were used for image acquisition. Phase contrast and widefield fluorescence microscopy images (minimum of five fields of view per mutant) were acquired with an exposure time of 100 ms for all channels.

### Analysis of cell morphology of NTML mutants

eHooke (18) was used to automatically perform image segmentation of cells from each mutant (at least five fields of view per mutant) and to measure morphological parameters of individual cells, namely area, perimeter, axis sizes, eccentricity and irregularity. This was followed by a visual inspection of the images by at least two users to qualitatively evaluate the quality of segmentation results. Data from images that were not correctly segmented was not considered for further analysis. Mutants were ranked by eccentricity values (which can vary between 0 and 1), calculated according to equation 1, where “a” and “b” are the semi-major and semi-minor axes, respectively, defined as the major and minor axes of the smallest rectangle that can contain each cell (18).

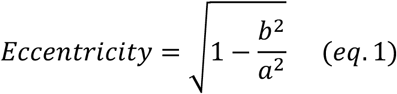

### Construction of plasmids and bacterial strains

The plasmids and bacterial strains used in this study are described in Tables 1 and 2, respectively. The primers used are listed in Table 3.

Deletion mutants were constructed using the thermosensitive plasmid pMAD (19). The upstream and downstream regions of the genes SAUSA300_1090, SAUSA300_2249 (*ssaA*), SAUSA300_0128, SAUSA300_1337 (*gpsB*), SAUSA300_1113 (*pknB*) and SAUSA300_1175 (*rodZ*) were amplified from *S. aureus* JE2 genomic DNA using the primers identified in Table 3. Subsequently, these fragments were cloned into pMAD plasmid in the SmaI site, in *E. coli* DC10B, using NEBuilder HiFi DNA assembly mix (New England BioLabs) through the Gibson assembly technique (20) creating pMAD Δ*1090*, pMAD Δ*ssaA*, pMAD Δ*0128*, pMAD Δ*pknB* and pMAD Δ*rodZ* (see Table 1). For pMAD Δ*gpsB*, upstream and downstream regions were amplified, joined by overlap PCR and cloned into pMAD using NcoI and BamHI restriction sites. Plasmids pMAD Δ*pbp3* (8), pMAD Δ*rodA* (8), pMAD Δ*mreD* (21), pMAD Δ*mreC* (21) and pMAD Δ*pbp4* (22) were already available.

The plasmids were extracted using a miniprep kit (NZYTech), the constructs were confirmed by Sanger sequencing (STAB VIDA) and then electroporated into *S. aureus* RN4220 at 30 °C using erythromycin and X-gal selection (23). Blue colonies were selected to produce a phage lysate using 80α (24) and the plasmids were transduced to *S. aureus* JE2. Constructs were integrated into the chromosome at 43 °C, as previously described (19). Cells were then incubated at 30 °C and colonies where the plasmid had undergone excision were selected, tested by PCR, resulting in strains JE2 Δ*1090*, JE2 Δ*ssaA*, JE2 Δ*pbp3*, JE2 Δ*rodA*, JE2 Δ*0128*, JE2 Δ*mreD*, JE2 Δ*gpsB*, JE2 Δ*pknB*, JE2 Δ*mreC*, JE2 Δ*rodZ*, JE2 Δ*pbp4*.

Using the same procedure, pMAD Δ*gpsB* vector was transduced into strains COL, ColsGFP-PBP1, BCBPM073, ColsGFP-PBP3, COLpPBP4-YFP (25) and COL EzrA-sGFP (26) to delete *gpsB*, resulting in strains COL Δ*gpsB*, ColsGFP-PBP1 Δ*gpsB*, ColsGFP-PBP2 Δ*gpsB*, ColsGFP-PBP3 Δ*gpsB*, COLpPBP4-YFP Δ*gpsB*, COL EzrA-sGFP Δ*gpsB*. Deletion of each gene was confirmed by PCR using primers in Table 3 followed by Sanger sequencing, across the deleted region.

To complement the deletion mutants with plasmid-encoded copies of the corresponding gene, each gene was amplified by PCR, using primers indicated in Table 3, and cloned into the pCNX plasmid under the control of P_*cad*_ promoter (7), using restriction enzymes SmaI and EcoRI for *mreC* and *mreD* fragments and NEBuilder HiFi DNA assembly mix (New England BioLabs) through the Gibson assembly technique (20) for the remaining fragments. The following plasmids were constructed: pCNX *1090*, pCNX *ssaA*, pCNX *0128*, pCNX *mreD*, pCNX *gpsB*, pCN51 *gpsB*, pCNX *pknB*, pCNX *mreC*, pCNX *rodZ* and pCNX *pbp4* (see Table 1). After confirmation of the insert sequence in pCNX by PCR and Sanger sequencing, DNA of plasmids with the correct insert was extracted and electroporated into *S. aureus* RN4220 at 37 °C using kanamycin selection (23). Each plasmid was then transduced into the corresponding deletion mutant using phage 80α, generating JE2 Δ*1090* pCNX *1090*, JE2 Δ*ssaA* pCNX *ssaA*, JE2 Δ*0128* pCNX *0128*, JE2 Δ*mreD* pCNX *mreD*, JE2 Δ*gpsB* pCNX *gpsB*, JE2 Δ*pknB* pCNX *pknB*, JE2 Δ*mreC* pCNX *mreC*, JE2 Δ*rodZ* pCNX *rodZ and* JE2 Δ*pbp4* pCNX *pbp4*. As a control, the pCNX empty vector was transduced into *S. aureus* JE2 strain, generating JE2 pCNX.

### Fluorescence microscopy and cell morphology image analysis

For imaging, *S. aureus* strains were grown overnight in TSB with selective antibiotics at 37°C, diluted to an OD_600nm_ of 0.05 and grown at 37 °C with selective antibiotics and appropriate inducers until OD_600nm_ of 0.5-0.6. When required, cells were labelled for 5 min at 37 °C, with agitation, with DNA dye Hoechst 33342 (1 μg/mL, Invitrogen) and membrane dye Nile Red (5 μg/mL, Invitrogen). The cells were centrifuged for 1 min at 10,000 rpm in a benchtop centrifuge (Eppendorf 5430), resuspended in PBS and 1 μL of this cell suspension was placed onto a gel pad composed of 1.2% agarose (TopVision, Thermo Fisher Scientific) prepared in PBS and mounted on a microscopy slide.

Images were acquired with a Zeiss Axio Observer Z1 microscope equipped with a Plan-Apochromat 100x/1.40 oil Ph3 objective with numerical aperture 0.55, HXP 120 V Illuminator (Zeiss) and Photometrics CoolSNAP HQ camera (Roper Scientific, Inc.) and controlled by a ZEN software (Zeiss). The filters Brightline TXRED-4040B and Brightline DAPI-1160A (Semrock) were used for image acquisition of cells labelled with Nile red and Hoechst 33342, respectively. Phase-contrast images and widefield microscopy images were acquired with 100 ms exposure time. At least 10 images per mutant were acquired, and three biological replicates were performed for each mutant.

Automated analysis of microscopy images was performed using eHooke (18) to measure morphological parameters including eccentricity, and to automatically assign the cell cycle phase to each analysed cell. Eccentricity was calculated using Equation 1 described above. Data was plotted using GraphPad Prism 8 (GraphPad Software).

### Assessment of FtsZ dynamics

To measure FtsZ treadmilling speed, COL EzrA-sGFP and COL EzrA-sGFP Δ*gpsB* strains were grown overnight in TSB and diluted 1:200 in fresh TSB followed by incubation with shaking at 37ºC. Exponentially growing cells (OD_600_ of 0.6-0.8) were harvested by centrifugation for 1 min at 9,300 g, resuspended in 30 μL fresh TSB, and spotted on a pad of 1.5% molecular biology grade agarose (Bio-Rad) in M9 minimal medium (KH_2_PO_4_ 3.4 g/L, VWR; K_2_HPO_4_ 2.9 g/L, VWR; di-ammonium citrate 0.7 g/L, Sigma-Aldrich; sodium acetate 0.26 g/L, Merck; glucose 1% (w/v), Merck; MgSO_4_ 0.7 mg/L, Sigma-Aldrich; CaCl_2_ 7 mg/L, Sigma-Aldrich; casamino acids 1% (w/v), Difco; MEM amino acids 1x, Thermo Fisher Scientific; MEM vitamins 1x, Thermo Fisher Scientific) mounted in a Gene Frame (Thermo Fisher Scientific) on a microscope slide. Imaging was performed in a DeltaVision OMX SR microscope equipped with a hardware-based focus stability (HW UltimateFocus) and an environmental control module set to 37 °C. Z-stacks of three images with a step size of 500 nm were acquired every three seconds for three minutes using a 488-nm laser (100 mW, at 10% maximal power) with an exposure time of 50 ms. Maximum intensity projection (MIP) of three images from each Z-stack and subsequent image deconvolution was performed for each time frame in the software SoftWoRx. All 61 time-frames were aligned using NanoJ-Core drift correction (27) and then used to perform MIP for the drawing of 1-pixel freehand lines over EzrA-sGFP in images of new born sister cells that are still attached via the septum of the mother cell, in which nascent Z-rings appear sparse and D-shaped. Space-time kymographs were generated by extracting fluorescence intensities from individual time frames along drawn freehand lines using the software Fiji (28). FtsZ treadmilling speed was calculated in nm/s by determining the slope of diagonals spanning the entire width of generated kymographs.

### Analysis of the localization of PBPs and of peptidoglycan synthesis

To localize fluorescent derivatives of PBPs 1-4, strains ColsGFP-PBP1, BCBPM073 (encoding sGFP-PBP2), ColsGFP-PBP3 and COLpPBP4-YFP, the corresponding *gpsB* deletion mutants ColsGFP-PBP1 Δ*gpsB*, ColsGFP-PBP2 Δ*gpsB*, ColsGFP-PBP3 Δ*gpsB*, COLpPBP4-YFP Δ*gpsB*, were grown in TSB 37 ºC to OD_600nm_ of 0.5. Images were acquired with the Zeiss Axio Observer Z1 microscope described above, with filters Brightline GFP-3035D and Brightline YFP-2427A to image GFP and YFP derivatives, respectively, with 4000 ms exposure time.

To evaluate localization of peptidoglycan synthesis activity, *S. aureus* cells of strain COL and COL Δ*gpsB* were grown in TSB at 37 ºC, in triplicate, to an OD_600nm_ of 0.4 and labelled with fluorescent D-amino-acid HADA (29) at 0.1mM for 30 min at 37 ºC, with agitation. Cells were then washed with PBS and placed on an 1.5% agarose pad mounted on a microscopy slide. Images were acquired with the Zeiss Axio Observer Z1 microscope described above using the filter Brightline DAPI-1160A (Semrock), with 100 ms exposure time.

For image analysis, phase 3 cells (with complete septum) were selected and the ratio of the fluorescence signal at the septum (considering only the 25% brightest pixels) versus the cell periphery was calculated using eHooke (18). Statistical analysis was performed using a two-sided Mann-Whitney U test, performed using the Python package SciPy version 1.10.1 (30).

## Results

### Screening of Nebraska Transposon Mutant Library (NTML) for mutants with reduced eccentricity

*S. aureus* cells undergo minor elongation during the cell cycle (7, 8). To the best of our knowledge, only two proteins, PBP3 and RodA (8), have been identified so far as being required for this elongation process. To identify new determinants required for *S. aureus* elongation, we labelled the 1920 NTML transposon insertion mutants in non-essential genes of JE2 (17), as well as the parental strain JE2, with a membrane dye and a DNA dye and imaged them by fluorescence microscopy. At least 5 fields of view were imaged for each strain, resulting in a library of over 10,000 images. Morphological parameters of each cell were automatically determined using eHooke software (18) and mutants were ranked from lowest (more spherical) to highest (more elongated) average cell eccentricity.

The top 7 mutants with lowest eccentricity were selected for further studies: SAUSA300_1090, *lgt, ssaA, pbp3, rodA*, SAUSA300_0128 and *mreD* (Table 4). The fact that 3^rd^ and 4^th^ positions in the ranking of increasing eccentricity were occupied by the genes encoding for RodA and PBP3, required for elongation (8), validates the screening. Repeating the imaging of selected transposon mutants did not confirm the decreased eccentricity of the *lgt* mutant, which was therefore considered a false positive from the screening and not studied further.

Of the top 100 hits from the eccentricity screening, we selected two further candidates, GpsB and PknB which we reasoned could have a role in elongation based on the function of these proteins in other organisms. GpsB contributes to the control of the cell elongation-division cycle in *Bacillus subtilis* by shuttling between the septum and the lateral wall, together with penicillin binding protein PBP1 (31). In *S. pneumoniae*, depletion of GpsB results in the formation of elongated, enlarged cells, presumably because GpsB mediates septal ring closure (32, 33). Given that the *B. subtilis* serine/threonine kinase PrkC phosphorylates GpsB (34), we also included in our study the main serine/threonine protein kinase present in *S. aureus*, PknB.

Two additional mutants lacking proteins with reported roles in elongation of other bacterial species, MreC and RodZ, were also included in the study. MreC, together with MreD, has a role in cell elongation through the coordination of peripheral PGN synthesis in *B. subtilis* and *S. pneumoniae* (14, 35). A previous study did not identify morphology changes in *S. aureus* mutants lacking MreD and MreC but their eccentricity was not determined (21). Given that the *mreD* mutant was a top hit in the screening, we decided to include also the *mreC* mutant in this study, although it ranked lower in the screening (Table 4). RodZ is a non-essential component of the elongasome (also known as Rod complex) of *E. coli* or *B. subtilis*, and *rodZ* deletion results in cell shape changes from rod to round or ovoid (36-38). Although a transposon mutant is absent from the NTML, we were able to generate a *rodZ* deletion mutant in JE2, showing that it is not an essential gene in this *S. aureus* strain.

Finally, we also included the NTML mutant lacking PBP4 because we previously showed that this protein is involved in peripheral peptidoglycan synthesis in *S. aureus* (7). We reasoned that it could potentially have a role in cell elongation via insertion of peripheral peptidoglycan, despite its low ranking in the screening for cell eccentricity (Table 4).

To confirm the results from the screening, and test if the morphology changes in the transposon mutants were not due to polar effects on downstream genes, new mutants were made by individually deleting each selected gene mentioned above from the genome of strain JE2. Furthermore, each deletion mutant was complemented with a plasmid expressing the corresponding gene. Deletion mutants and corresponding complemented strains were labelled with membrane and DNA dyes and imaged by widefield fluorescence microscopy. The eccentricity was measured only in cells with a closed septum (cell cycle phase 3 cells), since elongation is more pronounced in this phase (7, 8) (Figure 1). As expected, cells of all tested mutants had lower eccentricity (cells were more spherical) than cells of the parental strains JE2. However, for mutants in SAUSA300_1090, *ssaA*, SAUSA300_0128, *mreC, mreD* and *pbp4* this phenotype was not fully complemented by the corresponding plasmid-encoded gene, possibly due to the lack of native gene regulation or protein overexpression. The strongest phenotype, which was readily complemented by plasmid-encoded gene, was observed for cells of the *gpsB* deletion mutant, which was therefore selected for further studies.

**Figure 1:**
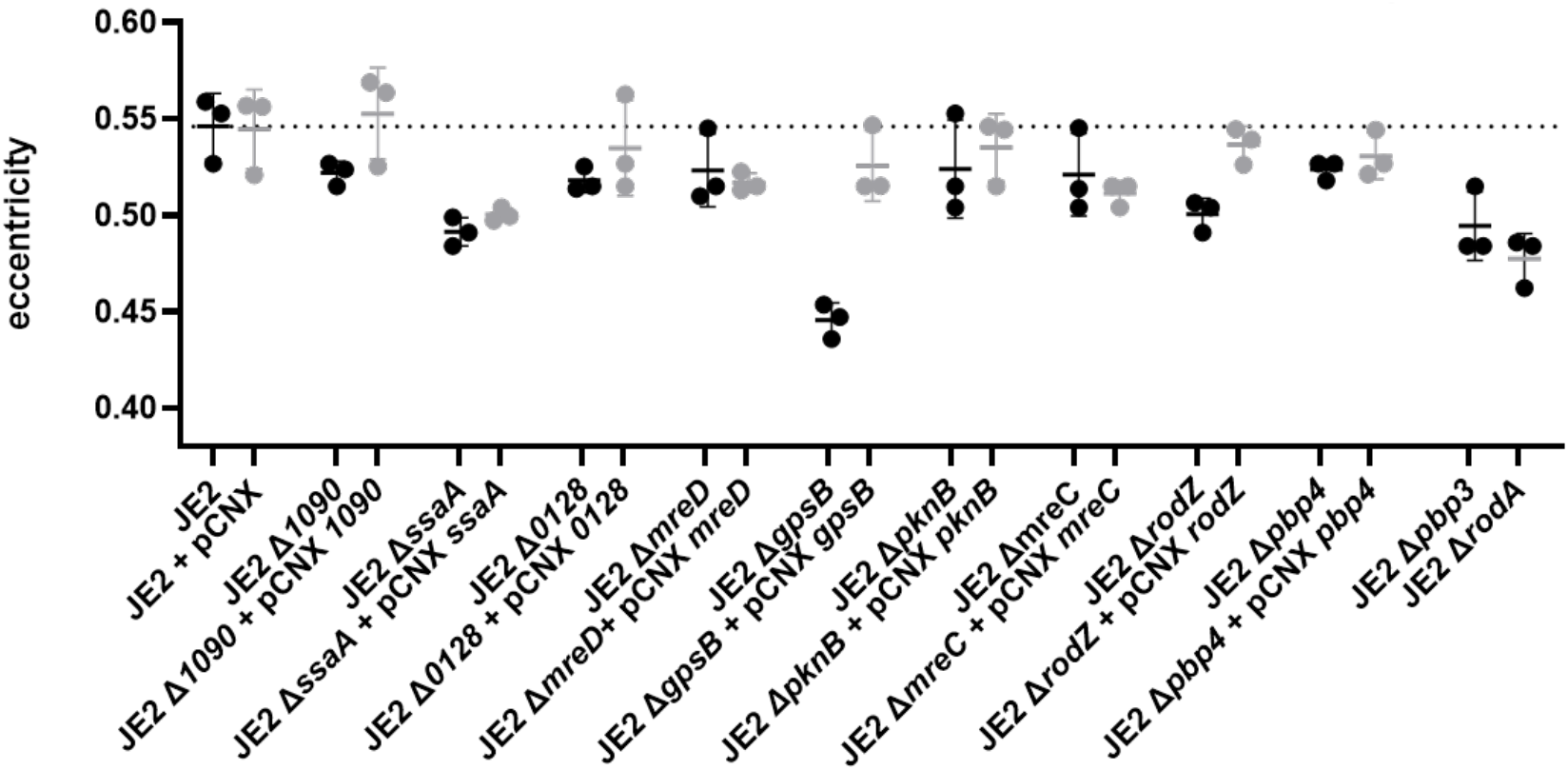
*S. aureus* mutants with reduced eccentricity. Cell eccentricity was measured in cells with a complete septum (cell cycle phase 3, n=550 cells/replicate, 3 replicates per strain) of selected mutants (black circles). Each mutant was complemented the corresponding gene encoded in pCNX replicative plasmid (grey circles). The empty vector was introduced in the parental strain JE2. Black lines represent the mean eccentricity and standard deviation for each strain. The average eccentricity of JE2 cells is indicated by the dashed line.

### Lack of GpsB does not alter FtsZ treadmilling speed

GpsB is a small coiled-coil protein, that localizes at midcell, previously reported to be essential in *S. aureus* strain SH1000 (39). However, a transposon mutant in *gpsB* is present in the NTML (17) and in additional reported transposon screenings (40) and a deletion mutant of *gpsB* has been recently reported (41). We could also easily delete *gpsB* from both COL and JE2 strains, indicating that *gpsB* is dispensable for growth in various *S. aureus* strains. The function of GpsB in *S. aureus* is still not fully elucidated, but it has been proposed to promote stabilization of the Z-ring at the onset of cell division (39). This would result in higher local concentration of FtsZ, activation of its GTPase activity and triggering of FtsZ treadmilling (39). In *S. aureus*, FtsZ treadmilling is essential during the early stages of cytokinesis (42). We hypothesized that regulation of FtsZ treadmilling activity by GpsB could modulate peptidoglycan synthesis at the septum, causing the morphology changes observed in the *gpsB* mutant. To measure FtsZ treadmilling speed, we used as a proxy a functional fusion of superfast GFP (sGFP) to EzrA, a direct interaction partner of FtsZ. We have previously shown that FtsZ and EzrA undergo similar movement dynamics, sensitive to FtsZ inhibitor PC190723 (42). Our quantitative analysis focused on the early stages of cytokinesis, in which nascent Z-rings appear sparse and D-shaped, in new born sister cells that are still attached via the septum of the mother cell (Figure 2a), given that GpsB was proposed to stabilize FtsZ at the onset of cell division (39). We observed that in TSB rich medium at 37°C, FtsZ filaments/bundles moved at the same speed in the parental strain (58.7±7.7 nm/s; *n*=43) and the *gpsB* deletion strain (59.6±7.6 nm/s; *n*=41) (Figure 2b), indicating that GpsB does not increase FtsZ treadmilling. Therefore *S. aureus* cell morphology changes in the *gpsB* mutant are not mediated by regulation of FtsZ treadmilling.

**Figure 2.**
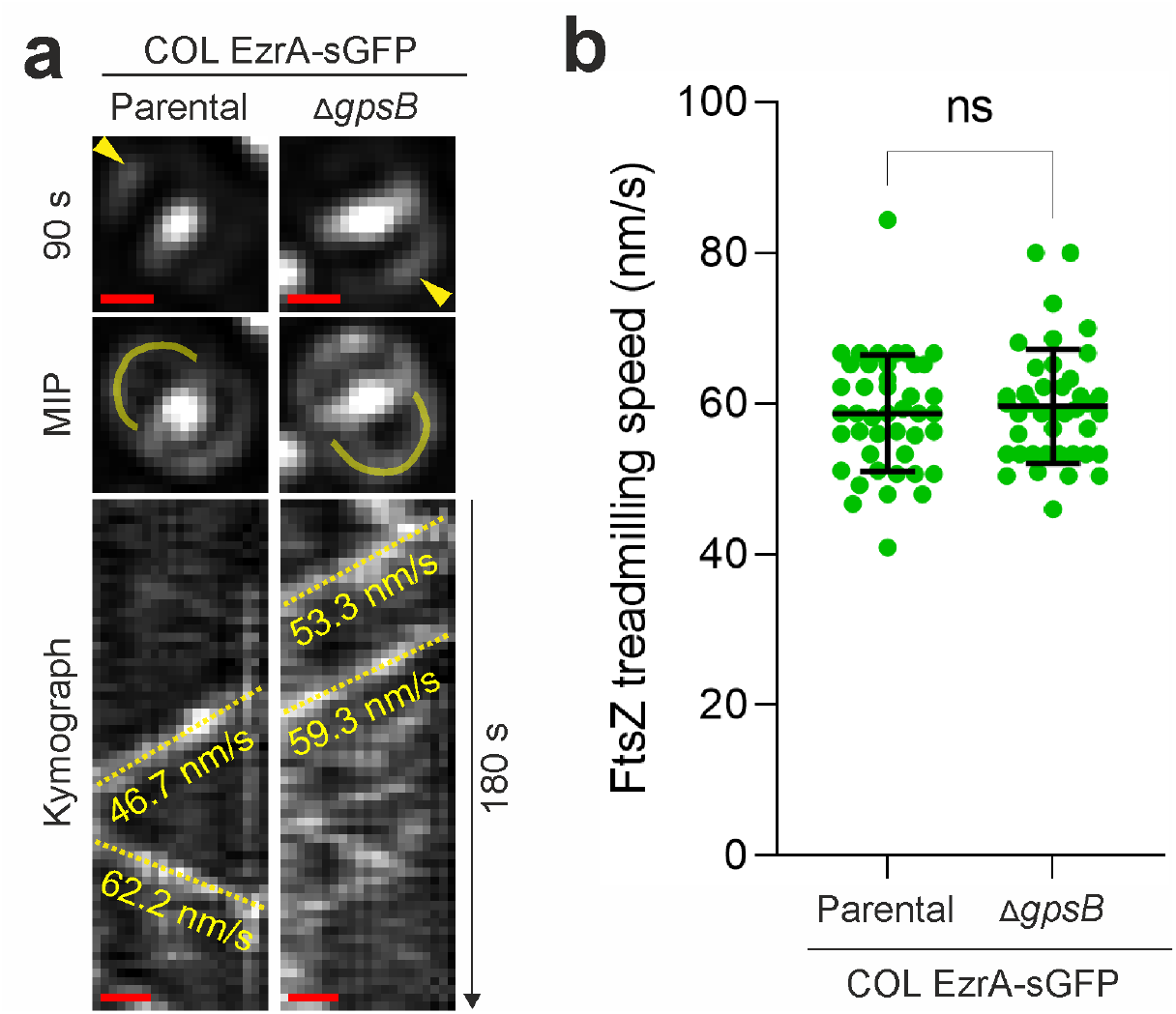
GpsB does not affect FtsZ treadmilling speed *in vivo*. **(a)** Representative epifluorescence images of EzrA-sGFP in pre-divisional cells with nascent Z-rings of strains COL EzrA-sGFP (left) and COL EzrA-sGFP Δ*gpsB* (right) at 90 s and throughout a 180-s time series (MIP, maximum intensity projection). Yellow arrow heads indicate an EzrA-sGFP patch whose change in localization was followed over time. Kymographs were generated by extracting fluorescence intensity values along indicated yellow lines. Yellow dashed lines in kymographs indicate the slopes used to calculate EzrA-sGFP movement, a proxy for FtsZ treadmilling. Scale bars, 0.5 μm. **(b**) FtsZ treadmilling speed measured in indicated genetic backgrounds. Data are represented as scatter plots in which the middle line represents the mean and the top and bottom lines show the standard deviation of slopes determined from diagonal lines in kymographs. n>30/sample, statistical analysis was performed using a two-tailed Mann–Whitney *U*-test (*U*=846; *P*=0.7531).

### PBP2 and PBP4 partially delocalize from the septum in the absence of GpsB

Besides interacting with FtsZ (39), *S. aureus* GpsB also interacts with PBP4 through a signature GpsB recognition sequence, suggesting that it may link cell division and peptidoglycan synthesis (43). If correct localization of one or more *S. aureus* PBPs was dependent on GpsB, then its absence could result in peptidoglycan incorporation in incorrect cellular locations, which could lead to morphological alterations. To test if this was the case, we deleted *gpsB* in four previously constructed strains expressing fluorescent fusions to each of the four *S. aureus* native PBPs, PBP1-4, and imaged the resulting strains by fluorescence microscopy. PBP1, an essential class b PBP with transpeptidase activity, was previously show to localize at the septum. This localization was not altered in the absence of GpsB, as in both ColsGFP-PBP1 and ColsGFP-PBP1 Δ*gpsB* the sGFP-PBP1 fluorescent signal was present at the septum and absent from the peripheral membrane (Figure 3a). We also did not observe any significant change in the localization pattern of PBP3 (Figure 3a). For strains expressing fluorescent derivatives of PBP2 and PBP4, we calculated the ratio of the fluorescence signal at the septum versus the cell periphery (Figure 3b). In both cases this ratio decreases in the absence of GpsB, indicating partial delocalization of PBP2 and PBP4 from the septum to the cell periphery. PBP2 is the only bifunctional PBP in *S. aureus*, with both transglycosylase and transpeptidase activity, for synthesis of the glycan strands and crosslinking via peptide bridges, respectively (44). PBP4 is a low molecular weight PBP with transpeptidase activity, responsible for the high levels of crosslinking characteristic of *S. aureus*, shown to be required for peptidoglycan synthesis at the cell periphery (7, 45).

**Figure 3.**
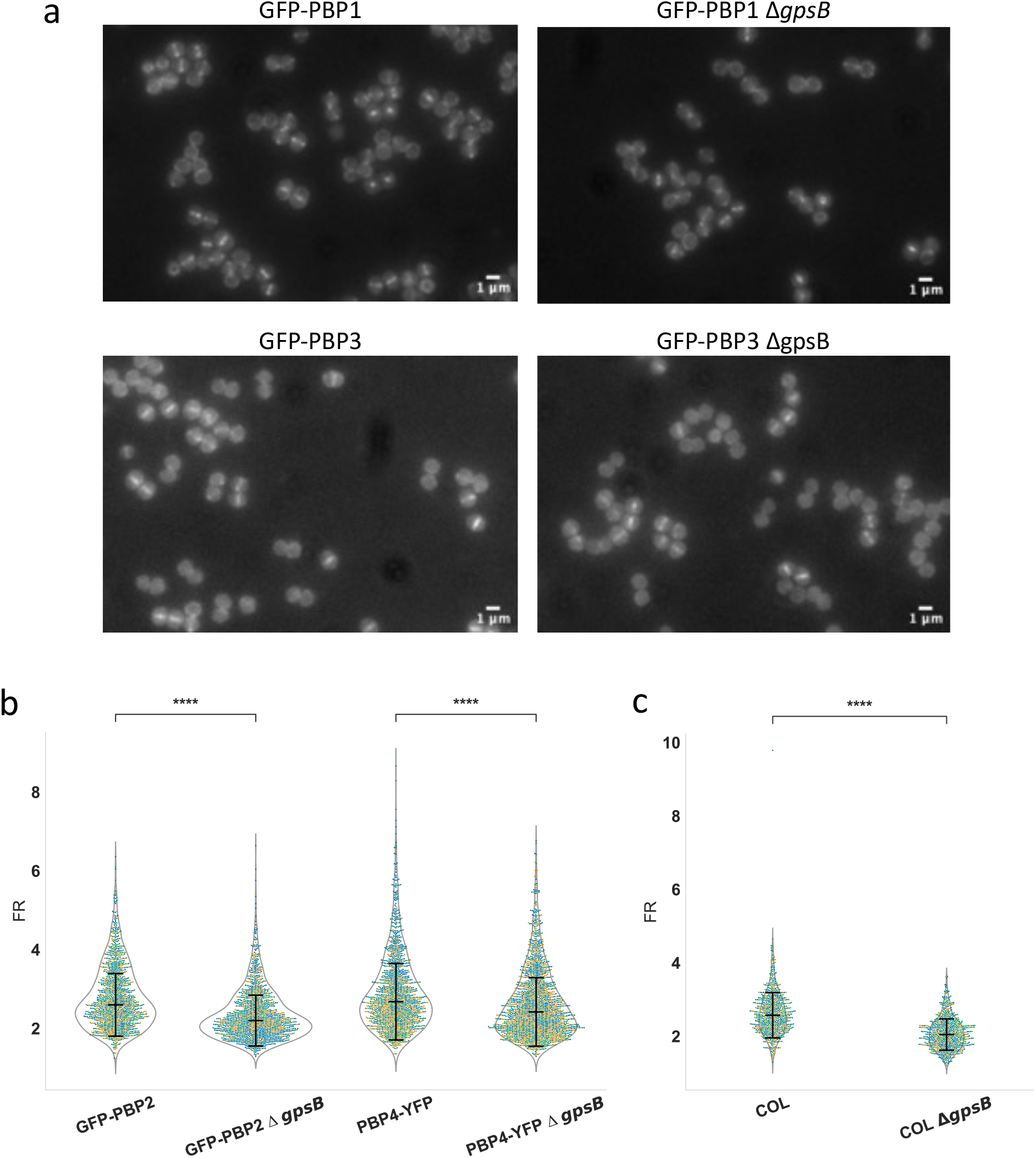
PBP2 and PBP4 partially delocalize from the septum in the absence of GpsB, leading to increased peptidoglycan synthesis at the cell periphery. Strains ColsGFP-PBP1, BCBPM073 (encoding sGFP-PBP2), ColsGFP-PBP3 and COLpPBP4-YFP, expressing fluorescent derivatives of the four *S. aureus* native PBPs, as well as the corresponding *gpsB* deletion mutants were imaged by fluorescence microscopy. **(a)** No change was observed in the localization pattern of the fluorescent derivatives of PBP1 and PBP3 in the absence of GpsB. **(b)** PBP2 and PBP4 partially delocalized from the division septum in the absence of GpsB. Graph shows the ratio of the fluorescent signal at the septum versus the peripheral membrane (FR), in cells with a complete septum, which decreases with protein delocalization from the septum. **(c)** *S. aureus* COL and COL Δ*gpsB* cells were labelled for 30 min with HADA, a fluorescent D-alanine is incorporated in peptidoglycan. Graph shows the ratio of the fluorescent signal at the septum versus the peripheral membrane (FR), in cells with a complete septum. This ratio is lower for COL Δ*gpsB* strain indicating that there is higher peptidoglycan synthesis activity at the cell periphery in the absence of GpsB. **(b**,**c)** Experiments were done in triplicate with yellow, green and blue dots corresponding to each replicate (n> 350 for GFP-PBP2 samples, n>450 for PBP4-YFP samples; n> 220 for COL; n>190 for COL Δ*gpsB*). Statistical analysis was performed using a two-sided Mann-Whitney U test, ****P<0.0001.

Partial delocalization of PBP2 and PBP4 to the cell periphery should lead to increased peptidoglycan synthesis over the entire cell surface. We labelled strains COL and COL Δ*gpsB* with HADA, a fluorescent derivative of 3-amino-D-alanine which is specifically incorporated in the pentapeptide chain of peptidoglycan (29), and observed that peptidoglycan synthesis at the septum versus the cell periphery decreases in the absence of GpsB (Figure 4). This increased peripheral synthesis may override the RodA/PBP3-mediated synthesis at the outer edge of the septum, previously shown to be required for *S. aureus* elongation (8), abolishing cell elongation in the *gpsB* mutants.

## Discussion

Elongation of *S. aureus* is dispensable *in vitro*, as mutants lacking the RodA/PBP3 pair of peptidoglycan synthetases, necessary for elongation, exhibit no growth defects in laboratory conditions such as TSB rich medium and 37ºC (8). However, the retention of an (albeit minor) elongation capacity through evolution, implies potential advantages in alternative conditions. Interestingly, in a murine model of osteomyelitis, an infection primarily caused by *S. aureus*, staphylococcal cells were observed as submicron rod-shaped bacteria within the canaliculi of live cortical bone (46). This suggests that deformation/elongation of *S. aureus* may be required for migration in bones during osteomyelitis (46). Furthermore, while wild type USA300 *S. aureus* cells were capable of propagation through a nanopore with a diameter of 0.5μm, smaller than the diameter of the cells, mutants lacking PBP3 or PBP4 showed reduced propagation through the nanopores, suggesting impaired ability to undergo the required deformation (47, 48). These mutants also failed to invade and colonize the osteocyte lacuno-canalicular network of cortical bone (47, 48), indicating that the ability to deform/elongate is required during osteomyelitis. We therefore think that understanding the mechanism of elongation of *S. aureus* is relevant not only for expanding our knowledge of cocci morphogenesis, but also for gaining a better understanding of how this bacterial pathogen causes an infection that is often regarded as incurable (46).

In this work we analysed the impact on the eccentricity of *S. aureus* cells, of deleting eleven non-essential genes. Two of these genes encoded RodA and PBP3, a SEDS/PBP pair previously reported to be involved in *S. aureus* elongation (8). The remaining nine mutants all had decreased eccentricity, indicating that elongation of *S. aureus* may be a complex process requiring multiple determinants. However, the phenotype of six of these mutants was very mild, including of mutants lacking two proteins of unknown function (SAUSA300_1090 and SAUSA300_0128), mutants lacking MreCD, two proteins with a well-studied role in elongation of rods (14, 49), the serine/threonine protein kinase PknB and PBP4, a peptidoglycan transpeptidase involved in peripheral peptidoglycan synthesis in *S. aureus* (7) (Figure 1). The remaining three mutants showed a stronger phenotype, with eccentricity being reduced by close to 10%. These include mutants lacking (i) RodZ, a member of the elongasome of rods in which ot interacts with MreC, MreD and RodA/bPBPs, possibly modulating MreB filament density in *B. subtilis* (16) and linking MreB filaments to the elongation peptidoglycan synthases (15). RodZ is also present in ovococcoid bacteria such as *S. pneumoniae*, where it seems to act as a scaffold protein of the elongasome despite the absence of MreB (50), a role that may be similar to that in *S. aureus*; (ii) SsaA, an autolysin with amidase activity (51). For peptidoglycan to expand in surface area, a process presumably required for cell elongation, both synthesis and hydrolysis are required, to allow for insertion of new material and relaxation of old material. SsaA may have a role in the relaxation of the peptidoglycan mesh at midcell during elongation, but further studies are required to prove this hypothesis; (iii) GpsB, a small membrane associated protein, widely conserved in Firmicutes, and proposed to coordinate peptidoglycan synthesis for cell growth and division, by binding cytoplasmic mini-domains of PBPs to ensure their correct subcellular localization (52). In *S. pneumoniae*, which lacks MreB, similarly to *S. aureus*, GpsB is part of a molecular switch that orchestrates peripheral and septal PG synthesis, and cells elongate in the absence of GpsB (33). The *S. aureus* mutant lacking GpsB was the only one in this study with cells more spherical (lower eccentricity) than those of the RodA/PBP3 mutants, pointing to an important role of this protein in staphylococcal elongation. We therefore focused on understanding the mechanism by which GpsB regulates elongation in *S. aureus*.

GpsB was previously suggested to regulate FtsZ treadmilling (39), which could alter peptidoglycan synthesis activity at the septum. However, we did not see any change in FtsZ treadmilling speed upon deletion of *gpsB*. Alternatively, given that GpsB was recently shown to interact with PBP4 in *S. aureus* (43), we hypothesized that GpsB could modulate *S. aureus* cell shape by directing PBPs to the cell periphery or to the septum.

In agreement with this hypothesis, we observed that both PBP2 and PBP4 partially delocalize from the septum to the peripheral wall in the absence of GpsB. This results in higher peptidoglycan synthesis activity in the cell periphery, which was confirmed by increased incorporation of the fluorescent D-amino acid HADA in the peptidoglycan of the peripheral wall. Localized peptidoglycan synthesis at the sidewall of the septal region is required for cell elongation in *S. aureus* (8). We propose that the increased peptidoglycan synthesis at the cell periphery in the absence of GpsB overrides the localized RodA/PBP3-mediated peptidoglycan synthesis at midcell, leading to more spherical cells. Conversely, in wild type cells that express GpsB, PBP2 and PBP4 activities are more restricted to the septum and expansion of the peripheral wall occurs mostly near the outer edge of the septum, allowing mild elongation of *S. aureus* cells. This role of GpsB does not necessarily imply that it is a member of an elongasome complex, but rather that it has a role in ensuring correct morphogenesis of *S. aureus* by contributing to the regulation of the spatial localization of PBPs.

## Supporting information

Slupplemental Tables

## Acknowledgements

We thank Zach Hensel (ITQB-NOVA) for helpful discussion and Mike VanNieuwenhze (Indiana University) for the generous gift of HADA. This study was funded by the European Research Council through grant ERC-2017-CoG-771709 (to M.G.P.), by national funds from FCT– Fundação para a Ciência e a Tecnologia through PTDC/BIA-MIC/6982/2020 (to HV), MOSTMICRO-ITQB R&D Unit (UIDB/04612/2020, UIDP/04612/2020 to ITQB-NOVA) and LS4FUTURE Associated Laboratory (LA/P/0087/2020 to ITQB-NOVA); contract 2022.03033.CEECIND (to SS), fellowships PD/BD/135480/2018 (to SFC), COVID/BD/152499/2022 (to SFC), SFRH/BD/147052/2019 (to BMS), UI/BD/153384/2022 (to LBM), SFRH/BD/143461/2019 (to AMD), 2021.06849.BD (to ADB) and by the European Union’s Horizon 2020 research and innovation programme under the Marie Sklodowska-Curie grant agreement No 839596 (to SS).

## BIBLIOGRAPHY

1. Murray CJL, Ikuta KS, Sharara F, Swetschinski L, Aguilar GR, Gray A, Han C, Bisignano C, Rao P, Wool E, Johnson SC, Browne AJ, Chipeta MG, Fell F, Hackett S, Haines-Woodhouse G, Hamadani BHK, Kumaran EAP, McManigal B, Agarwal R, Akech S, Albertson S, Amuasi J, Andrews J, Aravkin A, Ashley E, Bailey F, Baker S, Basnyat B, Bekker A, Bender R, Bethou A, Bielicki J, Boonkasidecha S, Bukosia J, Carvalheiro C, Castaneda-Orjuela C, Chansamouth V, Chaurasia S, Chiurchiu S, Chowdhury F, Cook AJ, Cooper B, Cressey TR, Criollo-Mora E, Cunningham M, Darboe S, Day NPJ, De Luca M, Dokova K, et al. 2022. Global burden of bacterial antimicrobial resistance in 2019: a systematic analysis. Lancet 399:629–655.

2. van Teeffelen S, Renner LD. 2018. Recent advances in understanding how rodlike bacteria stably maintain their cell shapes. F1000Res 7:241.

3. den Blaauwen T, Hamoen LW, Levin PA. 2017. The divisome at 25: the road ahead. Curr Opin Microbiol 36:85–94.

4. Chastanet A, Carballido-Lopez R. 2012. The actin-like MreB proteins in Bacillus subtilis: a new turn. Front Biosci (Schol Ed) 4:1582–1606.

5. Pinho MG, Kjos M, Veening J-W. 2013. How to get (a)round: mechanisms controlling growth and division of coccoid bacteria. Nat Rev Microbiol 11:601–614.

6. Lleo M. M. CP, Satta G. 1990. Bacterial cell shape regulation: testing of additional predictions unique to the two-competing-sites model for peptidoglycan assembly and isolation of conditional rod-shaped mutants from some wild-type cocci. J Bacteriol 172:3758–3771.

7. Monteiro JM, Fernandes PB, Vaz F, Pereira AR, Tavares AC, Ferreira MT, Pereira PM, Veiga H, Kuru E, VanNieuwenhze MS, Brun YV, Filipe SR, Pinho MG. 2015. Cell shape dynamics during the staphylococcal cell cycle. Nat Commun 6:8055.

8. Reichmann NT, Tavares AC, Saraiva BM, Jousselin A, Reed P, Pereira AR, Monteiro JM, Sobral RG, VanNieuwenhze MS, Fernandes F, Pinho MG. 2019. SEDS-bPBP pairs direct lateral and septal peptidoglycan synthesis in Staphylococcus aureus. Nat Microbiol 4:1368–1377.

9. Sham LT, Tsui HC, Land AD, Barendt SM, Winkler ME. 2012. Recent advances in pneumococcal peptidoglycan biosynthesis suggest new vaccine and antimicrobial targets. Curr Opin Microbiol 15:194–203.

10. Trouve J, Zapun A, Arthaud C, Durmort C, Di Guilmi AM, Soderstrom B, Pelletier A, Grangeasse C, Bourgeois D, Wong YS, Morlot C. 2021. Nanoscale dynamics of peptidoglycan assembly during the cell cycle of Streptococcus pneumoniae. Curr Biol 31:2844–2856 e6.

11. van Teeffelen S, Wang S, Furchtgott L, Huang KC, Wingreen NS, Shaevitz JW, Gitai Z. 2011. The bacterial actin MreB rotates, and rotation depends on cell-wall assembly. Proc Natl Acad Sci U S A 108:15822–7.

12. Domínguez-Escobar J, Chastanet A, Crevenna AH, Fromion V, Wedlich-Söldner R, Carballido-López R. 2011. Processive movement of MreB-associated cell wall biosynthetic complexes in bacteria. Science 333:225–228.

13. Garner EC, Bernard R, Wang W, Zhuang X, Rudner DZ, Mitchison T. 2011. Coupled, circumferential motions of the cell wall synthesis machinery and MreB filaments in B. subtilis. Science 333:222–225.

14. Leaver M, Errington J. 2005. Roles for MreC and MreD proteins in helical growth of the cylindrical cell wall in Bacillus subtilis. Mol Microbiol 57:1196–1209.

15. Ago R, Shiomi D. 2019. RodZ: a key-player in cell elongation and cell division in Escherichia coli. AIMS Microbiol 5:358–367.

16. Sun Y, Hurlimann S, Garner E. 2023. Growth rate is modulated by monitoring cell wall precursors in Bacillus subtilis. Nat Microbiol 8:469–480.

17. Fey PD, Endres JL, Yajjala VK, Widhelm TJ, Boissy RJ, Bose JL, Bayles KW. 2013. A genetic resource for rapid and comprehensive phenotype screening of nonessential Staphylococcus aureus genes. MBio 4:e00537–12.

18. Saraiva BM, Krippahl L, Filipe SR, Henriques R, Pinho MG. 2021. eHooke: A tool for automated image analysis of spherical bacteria based on cell cycle progression. Biol Imaging 1:e3.

19. Arnaud M, Chastanet A, Debarbouille M. 2004. New vector for efficient allelic replacement in naturally nontransformable, low-GC-content, gram-positive bacteria. Appl Environ Microbiol 70:6887–6891.

20. Gibson DG, Young L, Chuang RY, Venter JC, Hutchison CA, 3rd, Smith HO. 2009. Enzymatic assembly of DNA molecules up to several hundred kilobases. Nat Methods 6:343–345.

21. Tavares AC, Fernandes PB, Carballido-Lopez R, Pinho MG. 2015. MreC and MreD proteins are not required for growth of Staphylococcus aureus. PLoS One 10:e0140523.

22. Reed P, Atilano ML, Alves R, Hoiczyk E, Sher X, Reichmann NT, Pereira PM, Roemer T, Filipe SR, Pereira-Leal JB, Ligoxygakis P, Pinho MG. 2015. Staphylococcus aureus survives with a minimal peptidoglycan synthesis machine but sacrifices virulence and antibiotic resistance. PLoS Pathog 11:e1004891.

23. Veiga H, Pinho MG. 2009. Inactivation of the SauI Type I Restriction-Modification system is not sufficient to generate Staphylococcus aureus strains capable of efficiently accepting foreign DNA. Appl Environ Microbiol 75:3034–3038.

24. Oshida T, Tomasz A. 1992. Isolation and characterization of a Tn551-autolysis mutant of Staphylococcus aureus. J Bacteriol 174:4952–4959.

25. Loskill P, Pereira PM, Jung P, Bischoff M, Herrmann M, Pinho MG, Jacobs K. 2014. Reduction of the peptidoglycan crosslinking causes a decrease in stiffness of the Staphylococcus aureus cell envelope. Biophys J 107:1082–1089.

26. Saraiva BM, Sorg M, Pereira AR, Ferreira MJ, Caulat LC, Reichmann NT, Pinho MG. 2020. Reassessment of the distinctive geometry of Staphylococcus aureus cell division. Nat Commun 11:4097.

27. Laine RF, Tosheva KL, Gustafsson N, Gray RDM, Almada P, Albrecht D, Risa GT, Hurtig F, Lindas AC, Baum B, Mercer J, Leterrier C, Pereira PM, Culley S, Henriques R. 2019. NanoJ: a high-performance open-source super-resolution microscopy toolbox. J Phys D Appl Phys 52:163001.

28. Schindelin J, Arganda-Carreras I, Frise E, Kaynig V, Longair M, Pietzsch T, Preibisch S, Rueden C, Saalfeld S, Schmid B, Tinevez JY, White DJ, Hartenstein V, Eliceiri K, Tomancak P, Cardona A. 2012. Fiji: an open-source platform for biological-image analysis. Nat Methods 9:676–682.

29. Kuru E, Velocity Hughes H, Brown PJ, Hall E, Tekkam S, Cava F, de Pedro MA, Brun YV, VanNieuwenhze MS. 2012. In situ probing of newly synthesized peptidoglycan in live bacteria with fluorescent D-amino acids. Angew Chem Int Ed Engl 51:12519–12523.

30. Virtanen P, Gommers R, Oliphant TE, Haberland M, Reddy T, Cournapeau D, Burovski E, Peterson P, Weckesser W, Bright J, van der Walt SJ, Brett M, Wilson J, Millman KJ, Mayorov N, Nelson ARJ, Jones E, Kern R, Larson E, Carey CJ, Polat I, Feng Y, Moore EW, VanderPlas J, Laxalde D, Perktold J, Cimrman R, Henriksen I, Quintero EA, Harris CR, Archibald AM, Ribeiro AH, Pedregosa F, van Mulbregt P, SciPy C. 2020. SciPy 1.0: fundamental algorithms for scientific computing in Python. Nat Methods 17:261–272.

31. Claessen D, Emmins R, Hamoen LW, Daniel RA, Errington J, Edwards DH. 2008. Control of the cell elongation-division cycle by shuttling of PBP1 protein in Bacillus subtilis. Mol Microbiol 68:1029–1046.

32. Land AD, Tsui HC, Kocaoglu O, Vella SA, Shaw SL, Keen SK, Sham LT, Carlson EE, Winkler ME. 2013. Requirement of essential Pbp2x and GpsB for septal ring closure in Streptococcus pneumoniae D39. Mol Microbiol 90:939–955.

33. Fleurie A, Manuse S, Zhao C, Campo N, Cluzel C, Lavergne JP, Freton C, Combet C, Guiral S, Soufi B, Macek B, Kuru E, VanNieuwenhze MS, Brun YV, Di Guilmi AM, Claverys JP, Galinier A, Grangeasse C. 2014. Interplay of the serine/threonine-kinase StkP and the paralogs DivIVA and GpsB in pneumococcal cell elongation and division. PLoS Genet 10:e1004275.

34. Pompeo F, Foulquier E, Serrano B, Grangeasse C, Galinier A. 2015. Phosphorylation of the cell division protein GpsB regulates PrkC kinase activity through a negative feedback loop in Bacillus subtilis. Mol Microbiol 97:139–150.

35. Land AD, Winkler ME. 2011. The requirement for pneumococcal MreC and MreD is relieved by inactivation of the gene encoding PBP1a. J Bacteriol 193:4166–4179.

36. Alyahya SA, Alexander R, Costa T, Henriques AO, Emonet T, Jacobs-Wagner C. 2009. RodZ, a component of the bacterial core morphogenic apparatus. Proc Natl Acad Sci U S A 106:1239–44.

37. Bendezu FO, Hale CA, Bernhardt TG, Boer PAd. 2009. RodZ (YfgA) is required for proper assembly of the MreB actin cytoskeleton and cell shape in E. coli. EMBO Journal 28:193–204.

38. Muchova K, Chromikova Z, Barak I. 2013. Control of Bacillus subtilis cell shape by RodZ. Environ Microbiol 15:3259–3271.

39. Eswara PJ, Brzozowski RS, Viola MG, Graham G, Spanoudis C, Trebino C, Jha J, Aubee JI, Thompson KM, Camberg JL, Ramamurthi KS. 2018. An essential Staphylococcus aureus cell division protein directly regulates FtsZ dynamics. Elife 7.

40. Pang T, Wang X, Lim HC, Bernhardt TG, Rudner DZ. 2017. The nucleoid occlusion factor Noc controls DNA replication initiation in Staphylococcus aureus. PLoS Genet 13:e1006908.

41. Bartlett TM, Sisley TA, Mychack A, Walker S, Baker RW, Rudner DZ, Bernhardt TG. 2023. Identification of FacZ as a division site placement factor in Staphylococcus aureus. bioRxiv doi:10.1101/2023.04.24.538170.

42. Monteiro JM, Pereira AR, Reichmann NT, Saraiva BM, Fernandes PB, Veiga H, Tavares AC, Santos M, Ferreira MT, Macario V, VanNieuwenhze MS, Filipe SR, Pinho MG. 2018. Peptidoglycan synthesis drives an FtsZ-treadmilling-independent step of cytokinesis. Nature 554:528–532.

43. Sacco MD, Hammond LR, Noor RE, Bhattacharya D, Madsen JJ, Zhang X, Butler SG, Kemp MT, Jaskolka-Brown AC, Khan SJ, Gelis I, Eswara PJ, Chen Y. 2022. Staphylococcus aureus FtsZ and PBP4 bind to the conformationally dynamic N-terminal domain of GpsB. bioRxiv doi:10.1101/2022.10.25.513704.

44. Pinho M, de Lencastre H, Tomasz A. 2001. An acquired and a native penicillin-binding protein cooperate in building the cell wall of drug-resistant staphylococci. PNAS 98:10886–10891.

45. Memmi G, Filipe S, Pinho M, Fu Z, Cheung A. 2008. Staphylococcus aureus PBP4 is essential for Beta-lactam resistance in community-acquired methicillin-resistant strains. Antimicrob Agents Chemother 52:3955–3966.

46. de Mesy Bentley KL, Trombetta R, Nishitani K, Bello-Irizarry SN, Ninomiya M, Zhang L, Chung HL, McGrath JL, Daiss JL, Awad HA, Kates SL, Schwarz EM. 2017. Evidence of Staphylococcus aureus deformation, proliferation, and migration in canaliculi of live cortical bone in murine models of osteomyelitis. J Bone Miner Res 32:985–990.

47. Masters EA, Muthukrishnan G, Ho L, Gill AL, de Mesy Bentley KL, Galloway CA, McGrath JL, Awad HA, Gill SR, Schwarz EM. 2021. Staphylococcus aureus cell wall biosynthesis modulates bone invasion and osteomyelitis pathogenesis. Front Microbiol 12:723498.

48. Masters EA, de Mesy Bentley KL, Gill AL, Hao SP, Galloway CA, Salminen AT, Guy DR, McGrath JL, Awad HA, Gill SR, Schwarz EM. 2020. Identification of Penicillin Binding Protein 4 (PBP4) as a critical factor for Staphylococcus aureus bone invasion during osteomyelitis in mice. PLoS Pathog 16:e1008988.

49. Liu X, Biboy J, Consoli E, Vollmer W, den Blaauwen T. 2020. MreC and MreD balance the interaction between the elongasome proteins PBP2 and RodA. PLoS Genet 16:e1009276.

50. Lamanna MM, Manzoor I, Joseph M, Ye ZA, Benedet M, Zanardi A, Ren Z, Wang X, Massidda O, Tsui HT, Winkler ME. 2022. Roles of RodZ and class A PBP1b in the assembly and regulation of the peripheral peptidoglycan elongasome in ovoid-shaped cells of Streptococcus pneumoniae D39. Mol Microbiol 118:336–368.

51. Osipovitch DC, Therrien S, Griswold KE. 2015. Discovery of novel S. aureus autolysins and molecular engineering to enhance bacteriolytic activity. Appl Microbiol Biotechnol 99:6315–6326.

52. Cleverley RM, Rutter ZJ, Rismondo J, Corona F, Tsui HT, Alatawi FA, Daniel RA, Halbedel S, Massidda O, Winkler ME, Lewis RJ. 2019. The cell cycle regulator GpsB functions as cytosolic adaptor for multiple cell wall enzymes. Nat Commun 10:261.

