## Supplementary material for "The role of GpsB in cell morphogenesis of *Staphylococcus aureus*": Slupplemental Tables

**Table 1: Plasmids used in this work**

| Plasmid | Description | Reference |
|---|---|---|
| **pMAD** | *E. coli* /*S. aureus* shuttle vector with a thermosensitive origin of replication for Gram-positive bacteria, $Amp^R$, $Ery^R$, *lacZ* | (1) |
| **pMAD Δ*1090*** | pMAD with upstream and downstream regions of SAUSA300_1090 for constructing a null mutant, $Amp^R$, $Ery^R$, *lacZ* | This work |
| **pMAD Δ*ssaA*** | pMAD with upstream and downstream regions of SAUSA300_2249 (*ssaA*) for constructing a null mutant, $Amp^R$, $Ery^R$, *lacZ* | This work |
| **pMAD Δ*pbp3*** | pMAD with upstream and downstream regions of SAUSA300_1512 (*pbp3*) for constructing a null mutant, $Amp^R$, $Ery^R$, *lacZ* | (2) |
| **pMAD Δ*rodA*** | pMAD with upstream and downstream regions of SAUSA300_2040 (*rodA*) for constructing a null mutant, $Amp^R$, $Ery^R$, *lacZ* | (2) |
| **pMAD Δ*0128*** | pMAD with upstream and downstream regions of SAUSA300_0128 for constructing a null mutant, $Amp^R$, $Ery^R$, *lacZ* | This work |
| **pMAD Δ*mreD*** | pMAD with upstream and downstream regions of SAUSA300_1604 (*mreD*) for constructing a null mutant, $Amp^R$, $Ery^R$, *lacZ* | (3) |
| **pMAD Δ*gpsB*** | pMAD with upstream and downstream regions of SAUSA300_1337 (*gpsB*) for constructing a null mutant, $Amp^R$, $Ery^R$, *lacZ* | This work |
| **pMAD Δ*pknB*** | pMAD with upstream and downstream regions of SAUSA300_1113 (*pknB*) for constructing a null mutant, $Amp^R$, $Ery^R$, *lacZ* | This work |
| **pMAD Δ*mreC*** | pMAD with upstream and downstream regions of SAUSA300_1605 (*mreC*) for constructing a null mutant, $Amp^R$, $Ery^R$, *lacZ* | (3) |
| **pMAD Δ*rodZ*** | pMAD with upstream and downstream regions of SAUSA300_1175 (*rodZ*) for constructing a null mutant, $Amp^R$, $Ery^R$, *lacZ* | This work |
| **pMAD Δ*pbp4*** | pMAD with upstream and downstream regions of SAUSA300_0629 (*pbp4*) for constructing a null mutant, $Amp^R$, $Ery^R$, *lacZ* | (4) |
| **pCNX** | *E. coli* /*S. aureus* shuttle vector containing a cadmium inducible $P_{cad}$ promoter; $Amp^R$, $Kan^R$ | (5) |
| **pCN51** | *E. coli* /*S. aureus* shuttle vector containing a cadmium inducible $P_{cad}$ promoter; $Amp^R$, $Ery^R$ | (6) |
| **pCNX *1090*** | pCNX encoding SAUSA300_1090 under the control of $P_{cad}$ | This work |
| **pCNX *ssaA*** | pCNX encoding SAUSA300_2249 (*ssaA*) under the control of $P_{cad}$ | This work |
| **pCNX *0128*** | pCNX encoding SAUSA300_0128 under the control of $P_{cad}$ | This work |
| **pCNX *mreD*** | pCNX encoding SAUSA300_1604 (*mreD*) under the control of $P_{cad}$ | This work |

| | | |
|---|---|---|
| **pCNX *gpsB*** | pCNX encoding SAUSA300_1337 (*gpsB*) under the control of $P_{cad}$ | This work |
| **pCNX *pknB*** | pCNX encoding SAUSA300_1113 (*pknB*) under the control of $P_{cad}$ | This work |
| **pCNX *mreC*** | pCNX encoding SAUSA300_1605 (*mreC*) under the control of $P_{cad}$ | This work |
| **pCNX *rodZ*** | pCNX encoding SAUSA300_1175 (*rodZ*) under the control of $P_{cad}$ | This work |
| **pCNX *pbp4*** | pCNX encoding SAUSA300_0629 (*pbp4*) under the control of $P_{cad}$ | (7) |
| **pCN51 *gpsB*** | pCN51 encoding SAUSA300_1337 (*gpsB*) under the control of $P_{cad}$ | This work |

Abbreviations: $Amp^R$ – ampicillin resistance; $Ery^R$ – erythromycin resistance $Kan^R$ – Kanamycin resistance; cad – cadmium

**Table 2: Bacterial strains used in this work**

| Strain | Description | Reference |
|---|---|---|
| ***E. coli*** | | |
| **DC10B** | *dam+ dcm+ ∆hsdRMS endA1 recA1* | (8) |
| ***S. aureus*** | | |
| **RN4220** | Restriction deficient derivative of NCTC8325-4 | (9) |
| **JE2** | CA-MRSA | (10) |
| **COL** | HA-MRSA | (11) |
| **JE2 Δ*1090*** | JE2 SAUSA300_1090 deletion mutant | This work |
| **JE2 Δ*ssaA*** | JE2 *ssaA* deletion mutant | This work |
| **JE2 Δ*pbp3*** | JE2 *pbp3* deletion mutant | (2) |
| **JE2 Δ*rodA*** | JE2 *rodA* deletion mutant | This work |
| **JE2 Δ*0128*** | JE2 SAUSA300_0128 deletion mutant | This work |
| **JE2 Δ*mreD*** | JE2 *mreD* deletion mutant | This work |
| **JE2 Δ*gpsB*** | JE2 *gpsB* deletion mutant | This work |
| **JE2 Δ*pknB*** | JE2 *pknB* deletion mutant | This work |
| **JE2 Δ*mreC*** | JE2 *mreC* deletion mutant | This work |
| **JE2 Δ*rodZ*** | JE2 *rodZ* deletion mutant | This work |
| **JE2 Δ*pbp4*** | JE2 *pbp4* deletion mutant | This work |
| **JE2 pCNX** | JE2 transformed with pCNX empty vector; $Kan^R$ | This work |
| **JE2 Δ*1090* pCNX *1090*** | JE2 SAUSA300_1090 deletion mutant transformed with pCNX *1090*; $Kan^R$ | This work |
| **JE2 Δ*ssaA* pCNX *ssaA*** | JE2 *ssaA* deletion mutant transformed with pCNX *ssaA*; $Kan^R$ | This work |
| **JE2 Δ*0128* pCNX *0128*** | JE2 SAUSA300_0128 deletion mutant transformed with pCNX *0128*; $Kan^R$ | This work |
| **JE2 Δ*mreD* pCNX *mreD*** | JE2 *mreD* deletion mutant transformed with pCNX *mreD*; $Kan^R$ | This work |
| **JE2 Δ*gpsB* pCNX *gpsB*** | JE2 *gpsB* deletion mutant transformed with pCNX *gpsB*; $Kan^R$ | This work |

| | | |
|---|---|---|
| **JE2 Δ*pknB* pCNX *pknB*** | JE2 *pknB* deletion mutant transformed with pCNX *pknB*; $Kan^R$ | This work |
| **JE2 Δ*mreC* pCNX *mreC*** | JE2 *mreC* deletion mutant transformed with pCNX *mreC*; $Kan^R$ | This work |
| **JE2 Δ*rodZ* pCNX *rodZ*** | JE2 *rodZ* deletion mutant transformed with pCNX *rodZ*; $Kan^R$ | This work |
| **JE2 Δ*pbp4* pCNX *pbp4*** | JE2 *pbp4* deletion mutant transformed with pCNX *pbp4*; $Kan^R$ | This work |
| **JE2 *lgt*::ΦΝΣ** | JE2 with a transposon inserted in the *lgt* gene (Nebraska transposon mutant library); $Ery^R$ | (10) |
| **ColsGFP-PBP1** | COL *pbpA::sgfp-pbpA* | (12) |
| **BCBPM073** | COL *pbpB::sgfp-pbpB* | (13) |
| **ColsGFP-PBP3** | COL *pbpC::sgfp-pbpC* | (12) |
| **COLpPBP4-YFP** | COL *pbpD::pbpD-yfp;* $Kan^R$ | (14) |
| **COL EzrA-sGFP** | COL *ezrA::ezrA-sgfp* | (15) |
| **COL Δ*gpsB*** | COL *gpsB* deletion mutant | This work |
| **ColsGFP-PBP1 Δ*gpsB*** | COL *pbpA::sgfp-pbpA ΔgpsB* | This work |
| **ColsGFP-PBP2 Δ*gpsB*** | COL *pbpB::sgfp-pbpB ΔgpsB* obtained by deleting *gpsB* in BCBPM073 | This work |
| **ColsGFP-PBP3 Δ*gpsB*** | COL *pbpC::sgfp-pbpC ΔgpsB* | This work |
| **COLpPBP4-YFP Δ*gpsB*** | COL *pbpD::pbpD-yfp; ΔgpsB* $Kan^R$ | This work |
| **COL EzrA-sGFP Δ*gpsB*** | COL *ezrA::ezrA-sgfp; ΔgpsB* | This work |

Abbreviations: $Amp^R$ – ampicillin resistance; $Ery^R$ – erythromycin resistance $Kan^R$ – Kanamycin resistance; cad – cadmium

**Table 3: Primers used in this work**

| Primer | Used for | Insert length (bp) | Sequence 5'-3' |
|---|---|---|---|
| pMAD_2_Up_ 1090_fwd | Amplification of upstream and downstream regions of 1090 gene to clone into pMAD plasmid | 996 | CGATGCATGCCATGGTACCCCAATTAAGTGTAGACGATTC |
| Up_down_1090_rev | | | GCACGACACAATTACTTAACCTCCTTCTCC |
| Up_down_1090_fwd | | 1000 | GGTTAAGTAATTGTGTCGTGCTATAATTACG |
| down1090_ pMAD_rev | | | CTTCTAGAATTCGAGCTCCCCAAGCATATGAAAACTTATTTATCATTC |
| pMAD_up_ ssaA_fwd | Amplification of upstream and downstream regions of *ssaA* gene to clone into pMAD plasmid | 1000 | CGATGCATGCCATGGTACCCCAACACACATGTAATTAATAATCTTATC |
| up_down_ssaA_rev | | | CATAGCCATCACTAATTTAAAAATATCCTCCTAAAAATTTTAAATC |
| up_down_ssaA_fwd | | 1000 | GGAGGATATTTTTAAATTAGTGATGGCTATGTTTACGC |
| down_ssaA_ pMAD | | | CTTCTAGAATTCGAGCTCCCGACAGATGGTTTCAAAGC |
| pMAD_up_0128_fwd2 | Amplification of upstream and downstream regions of *0128* gene to clone into pMAD plasmid | 937 | GATGCATGCCATGGTACCCCGCAAGTTATAGTGGTTGTTG |
| up_down_0128_rev2 | | | CTCGATTAGTTTGTCACCTTAAATATTCTCTTTATTCTTTAGTG |
| up_down_0128_fwd | | 1000 | GGGAATAGACTGACAAACTAATCGAGAGAC |
| down_0128_ pMAD_rev | | | CTTCTAGAATTCGAGCTCCCAGTGGATGATTATTAACGG |
| upGpsB P1N | Amplification of upstream and downstream regions of *gpsB* gene to clone into pMAD plasmid | 621 | CTTACCATGGGGCGTAACAAAAGAGGGTAC |
| downGpsB P3 | | | CTTTTGTATTTAGTAATTACATTTTTTCCACCTCATTAGAAAC |
| upGpsB P2 | | 973 | GTTTCTAATGAGGTGGAAAAAATGTAATTACTAAATACAAAAG |
| downGpsB P2 B | | | CGAGGATCCGATTTAACGCTTCTACCTTG |
| pMAD_dPknB_Up_Fwd | Amplification of upstream and downstream regions of *pknB* gene to clone into pMAD plasmid | 805 | CTATCGATGCATGCCATGGTACCCGTAAAGACAAATGCTAGAGG |
| dPknB_Up_Rev | | | TTATACATCATCATATATTTTACCTATCATACTTTATCACCTTCAATAGCCGCGA |
| dPknB_Down_Fwd | | 773 | ATGATAGGTAAAATATATGATGATGTATAAATATAATTGAAGTAAATGTACCGA |
| pMAD_dPknB_Down_Rev | | | GAAGCTTCTAGAATTCGAGCTCCCCCACTAAGTACTATAAGTCCAG |
| RodZ_P1_EcoRI | Amplification of upstream and downstream regions of *rodZ* gene to clone into pMAD plasmid | 850 | GCTGAATTCCACCATGACAGTTGCAGAG |
| RodZ_P2 | | | ATTTCTGTTATTCATAGCCTCCTTACACTTAC |
| RodZ_P3 | | 841 | GGAGGCTATGAATAACAGAAATAAATTAGTGAG |
| RodZ_P4_NcoI | | | TGCATTCCATGGTTATATCCCCCTTGTATCG |
| pMADI | Verification of insert in pMAD plasmid for sequencing | variable | CTCCTCCGTAACAAATTGAGG |
| pMADII | | | CGTCATCTACCTGCCTGGAC |

| | | | |
|---|---|---|---|
| pCNseqUPfwd | Verification of insert under $P_{cad}$ promoter for sequencing | variable | CATATCAGGCAGATAATG |
| pCNseqDWRev | | | CAAAATTATACATGTCAACG |
| pCNX_rbs_ 1090_fwd | Amplification of *1090* gene to clone in pCNX plasmid | 981 | GTCGACTCTAGAGGATCCCCAATATCTAAGGAGGTAATATAATGGAGACTTATGAATTTAAC |
| 1090_pCNX_ rev | | | CCTGAATTCGAGCTCGGTACCCTTATGCATCTCTTTTTCGAATTG |
| pCNX_rbs_ ssaA_fwd | Amplification of *ssaA* gene to clone in pCNX plasmid | 866 | GTCGACTCTAGAGGATCCCCAATATCTAAGGAGGTAATATAATGAAGAAAATCGCTACAGC |
| ssaA_pCNX_ rev | | | CTGAATTCGAGCTCGGTACCCTTAGTGAATGAAGTTATAACCAG |
| pCNX_rbs_ 0128_fwd | Amplification of *0128* gene to clone in pCNX plasmid | 687 | GTCGACTCTAGAGGATCCCCATATCTAAGGAGGTAATATATGGAAAAAAATGTAGAAAAATCATTC |
| 0128_pCNX_ rev | | | CTGAATTCGAGCTCGGTACCCTCAATCTTTTTTCGAGACATGG |
| pCNX_rbs_ gpsB_fwd | Amplification of *gpsB* gene to clone in pCNX plasmid | 408 | GTCGACTCTAGAGGATCCCCAATATCTAAGGAGGTAATATAATGTCAGATGTTTCATTG |
| gpsB_pCNX_ rev | | | CTGAATTCGAGCTCGGTACCCTTATTTACCAAATACAGCTTTTTCTAAG |
| pCNX_PknB_Up_Fwd | Amplification of *pknB* gene to clone in pCNX plasmid | 2055 | GTCGACTCTAGAGGATCCCCGGCTATTGAAGGTGATAAAGTATGATAGG |
| pCNX_PknB_Down_Rev | | | AATTCGAGCTCGGTACCCTTATACATCATCATAGCTGAC |
| pMreC_SmaI_P1 | Amplification of *mreC* gene to clone in pCNX plasmid | 881 | TCCCCCGGGGACATAATAGAGGTGTTCTG |
| BTH_mreC_EcoRI_P2 | | | CGGAATTCTTATTTATCCCTGCTTTC |
| pMreD_SmaI_P1 | Amplification of *rmreD* gene to clone in pCNX plasmid | 568 | TCCCCCGGGGATGAAAGCAGGGATAAATAATG |
| BTH_mreD_Nfus_EcoRI_P2 | | | CGGAATTCTTACCATTGACGACGTTTC |
| pCNX_rbs_ rodZ_fwd | Amplification of *rodZ* gene to clone in pCNX plasmid | 455 | GTCGACTCTAGAGGATCCCCAATATCTAAGGAGGTAATATATTGAAAACGGTCGGTGAAG |
| rodZ_pCNX_ rev | | | CTGAATTCGAGCTCGGTACCCTTAAAATATTAAAACTAACATGATCCATAAC |
| up_1090_seq_ fwd | Verification of *1090* gene deletion | 2166 | CGAACTGACATTCGAGTG |
| down_1090_ seq_rev | | | GACAAGTTGCTACAAGTC |
| up_ssaA_seq_ fwd | Verification of *ssaA* gene deletion | 2110 | GACGACTACCATCGTATG |
| down_ssaA_ seq_rev | | | GTTGCGATAGTAGCTGTAG |
| up_0128_Seq Exc_fwd | Verification of *0128* gene deletion | 830 | CTGGAGGATCTGATTACG |
| down_0128_ Seq Exc_rev | | | CGATTGGTGTTGTTGGTC |
| up_mreD_Seq Exc_fwd | Verification of *mreD* gene deletion | 451 | CAAGTGGATTAGCTGATC |
| down_mreD_ SeqExc_rev | | | GTATCTCCTTCGTTTACGTC |
| up_gpsB_Seq Exc_fwd | Verification of *gpsB* gene deletion | 547 | CTAGTCACACTGGTACTC |

| | | | |
|---|---|---|---|
| up_gpsB_SeqExc_fwd | | | CGAGTCCTAAGTTCTTCAAG |
| up_mreC_SeqExc_fwd | Verification of *mreC* gene deletion | 440 | CACGTTGGAGACAACTTC |
| down_mreC_SeqExc_rev | | | GCTGAGCAATAATGATACG |
| dPknB_Conf_Fwd | Verification of *pknB* gene deletion | 1914 | GAATGACCAACTAGAACATGC |
| dPknB_Conf_Rev | | | ACTGAATCCAGGTGTGTCT |
| up_rodZ_SeqExc_fwd | Verification of *rodZ* gene deletion | 1864 | CTAGAAGTGTTACTGGAAC |
| down_rodZ_SeqExc_rev | | | GCAATAATGGCAATTGAC |

**Table 4: Mutants of Nebraska Transposon Mutant Library ranked by increasing eccentricity**

| Rank. | Name | Gene Description | Accession # | Area* ($\mu m^2$) | Eccen. | Phase (%) | | |
|---|---|---|---|---|---|---|---|---|
| | | | | | | 1 | 2 | 3 |
| - | JE2 WT | - | - | 1.36 | 0.50214 | 41 | 25 | 34 |
| 1 | SAUSA300_ 1090 | hypothetical protein | SAUSA300_1090 | 1.17 | 0.43678 | 53 | 25 | 20 |
| 2 | *lgt* | prolipoprotein diacylglyceryl transferase | SAUSA300_0744 | 1.23 | 0.44113 | 47 | 24 | 27 |
| 3 | *ssaA* | secretor antigen precursor SsaA | SAUSA300_2249 | 1.25 | 0.44402 | 51 | 24 | 24 |
| 4 | *pbp3* | penicillin-binding protein 3 | SAUSA300_1512 | 1.28 | 0.44598 | 50 | 21 | 27 |
| 5 | *rodA* | rod shape-determining protein RodA | SAUSA300_2040 | 1.29 | 0.44654 | 52 | 16 | 30 |
| 6 | SAUSA300_0128 | hypothetical protein | SAUSA300_0128 | 1.13 | 0.44659 | 52 | 27 | 19 |
| 6 | *mreD* | rod shape-determining protein MreD | SAUSA300_1604 | 1.21 | 0.44659 | 41 | 22 | 35 |
| 67 | *gpsB* | cell cycle protein | SAUSA300_1337 | 1.27 | 0.44892 | 41 | 23 | 34 |
| 74 | *pknB* | protein kinase | SAUSA300_1113 | 1.21 | 0.45189 | 44 | 22 | 33 |
| 1150 | *mreC* | rod shape-determining protein MreC | SAUSA300_1605 | 1.16 | 0.48233 | 47 | 27 | 25 |
| 1532 | *pbp4* | penicillin-binding protein 4 | SAUSA300_0629 | 1.36 | 0.49382 | 48 | 20 | 30 |

Rank – ranking in the eccentricity screening; *Mean area of phase contrast images of cells, in $\mu m^2$; Eccen. – median eccentricity.

**BIBLIOGRAPHY**


1. Arnaud M, Chastanet A, Debarbouille M. 2004. New vector for efficient allelic replacement in naturally nontransformable, low-GC-content, gram-positive bacteria. Appl Environ Microbiol 70:6887-6891.

2. Reichmann NT, Tavares AC, Saraiva BM, Jousselin A, Reed P, Pereira AR, Monteiro JM, Sobral RG, VanNieuwenhze MS, Fernandes F, Pinho MG. 2019. SEDS-bPBP pairs direct lateral and septal peptidoglycan synthesis in *Staphylococcus aureus*. Nat Microbiol 4:1368-1377.

3. Tavares AC, Fernandes PB, Carballido-Lopez R, Pinho MG. 2015. MreC and MreD proteins are not required for growth of *Staphylococcus aureus*. PLoS One 10:e0140523.

4. Reed P, Atilano ML, Alves R, Hoiczyk E, Sher X, Reichmann NT, Pereira PM, Roemer T, Filipe SR, Pereira-Leal JB, Ligoxygakis P, Pinho MG. 2015. *Staphylococcus aureus* survives with a minimal peptidoglycan synthesis machine but sacrifices virulence and antibiotic resistance. PLoS Pathog 11:e1004891.

5. Monteiro JM, Fernandes PB, Vaz F, Pereira AR, Tavares AC, Ferreira MT, Pereira PM, Veiga H, Kuru E, VanNieuwenhze MS, Brun YV, Filipe SR, Pinho MG. 2015. Cell shape dynamics during the staphylococcal cell cycle. Nat Commun 6:8055.

6. Charpentier E, Anton AI, Barry P, Alfonso B, Fang Y, Novick RP. 2004. Novel cassette-based shuttle vector system for gram-positive bacteria. Appl Environ Microbiol 70:6076-6085.

7. Pereira PM. 2013. Peptidoglycan Assembly Machines: The *Staphylococcus aureus* Penicillin-Binding ProteinsUniversidade Nova de Lisboa.

8. Monk IR, Shah IM, Xu M, Tan MW, Foster TJ. 2012. Transforming the untransformable: application of direct transformation to manipulate genetically *Staphylococcus aureus* and *Staphylococcus epidermidis*. MBio 3:e00277-11.

9. Nair D, Memmi G, Hernandez D, Bard J, Beaume M, Gill S, Francois P, Cheung AL. 2011. Whole-genome sequencing of *Staphylococcus aureus* strain RN4220, a key laboratory strain used in virulence research, identifies mutations that affect not only virulence factors but also the fitness of the strain. J Bacteriol 193:2332-2335.

10. Fey PD, Endres JL, Yajjala VK, Widhelm TJ, Boissy RJ, Bose JL, Bayles KW. 2013. A genetic resource for rapid and comprehensive phenotype screening of nonessential *Staphylococcus aureus* genes. MBio 4:e00537-12.

11. Gill SR, Fouts DE, Archer GL, Mongodin EF, Deboy RT, Ravel J, Paulsen IT, Kolonay JF, Brinkac L, Beanan M, Dodson RJ, Daugherty SC, Madupu R, Angiuoli SV, Durkin AS, Haft DH, Vamathevan J, Khouri H, Utterback T, Lee C, Dimitrov G, Jiang L, Qin H, Weidman J, Tran K, Kang K, Hance IR, Nelson KE, Fraser CM. 2005. Insights on evolution of virulence and resistance from the complete genome analysis of an early methicillin-resistant *Staphylococcus aureus* strain and a biofilm-producing methicillin-resistant *Staphylococcus epidermidis* strain. J Bacteriol 187:2426-2438.

12. Monteiro JM, Pereira AR, Reichmann NT, Saraiva BM, Fernandes PB, Veiga H, Tavares AC, Santos M, Ferreira MT, Macario V, VanNieuwenhze MS, Filipe SR, Pinho MG. 2018. Peptidoglycan synthesis drives an FtsZ-treadmilling-independent step of cytokinesis. Nature 554:528-532.

13. Tan CM, Therien AG, Lu J, Lee SH, Caron A, Gill CJ, Lebeau-Jacob C, Benton-Perdomo L, Monteiro JM, Pereira PM, Elsen NL, Wu J, Deschamps K, Petcu M, Wong S, Daigneault E, Kramer S, Liang L, Maxwell E, Claveau D, Vaillancourt J, Skorey K, Tam J, Wang H, Meredith TC, Sillaots S, Wang-Jarantow L, Ramtohul Y, Langlois E, Landry F, Reid JC, Parthasarathy G, Sharma S, Baryshnikova A, Lumb KJ, Pinho MG, Soisson SM, Roemer T. 2012. Restoring methicillin-resistant *Staphylococcus aureus* susceptibility to β-lactam antibiotics. Sci Transl Med 4:126ra35.

14. Loskill P, Pereira PM, Jung P, Bischoff M, Herrmann M, Pinho MG, Jacobs K. 2014. Reduction of the peptidoglycan crosslinking causes a decrease in stiffness of the *Staphylococcus aureus* cell envelope. Biophys J 107:1082-1089.

15. Saraiva BM, Sorg M, Pereira AR, Ferreira MJ, Caulat LC, Reichmann NT, Pinho MG. 2020. Reassessment of the distinctive geometry of *Staphylococcus aureus* cell division. Nat Commun 11:4097.
